# EGR1 regulates oral epithelial cell responses to *Candida albicans* via the EGFR- ERK1/2 pathway

**DOI:** 10.1101/2023.03.31.535186

**Authors:** Ruth E. Dickenson, Aize Pellon, Nicole O. Ponde, Olivia Hepworth, Lydia F. Daniels Gatward, Julian R. Naglik, David L. Moyes

## Abstract

*Candida albicans* is a fungal pathobiont colonising mucosal surfaces of the human body, including the oral cavity. Under certain predisposing conditions, *C. albicans* invades mucosal tissues activating EGFR-MAPK signalling pathways in epithelial cells via the action of its peptide toxin candidalysin. However, our knowledge of the epithelial mechanisms involved during *C. albicans* colonisation is rudimentary. Here, we describe the role of the transcription factor early growth response protein 1 (EGR1) in human oral epithelial cells (OECs) in response to *C. albicans*. EGR1 expression increases in OECs when exposed to *C. albicans* independently of fungal viability, morphology, or candidalysin release, suggesting EGR1 is involved in the fundamental recognition of *C. albicans*, rather than in response to invasion or ‘pathogenesis’. Upregulation of EGR1 is mediated by EGFR via Raf1, ERK1/2 and NF-κB signalling but not PI3K/mTOR signalling. Notably, EGR1 mRNA silencing impacts on anti-*C. albicans* immunity, reducing GM-CSF, IL-1α and IL-1β release, and increasing IL-6 and IL-8 production. These findings identify an important role for EGR1 in priming epithelial cells to respond to subsequent invasive infection by *C. albicans* and elucidate the regulation circuit of this transcription factor after contact.

## INTRODUCTION

Fungal microbes are an essential part of the human microbiota at many body sites, contributing to both health and disease (1). Fungal infections are an increasing global health issue, most commonly manifesting as superficial mucosal infections. These number in the millions each year and, despite not representing a major threat to life, significantly decrease patient life quality (2). Superficial infections can lead to far more serious invasive infections with significant mortality rates that result in over a million annual deaths globally. The fungal pathogen *Candida albicans* colonises ≈70% of the healthy population, mainly in the oral cavity, gut and female genital tract (3, 4). However, under certain environmental conditions, this fungus can become pathogenic and cause superficial mucosal infections in otherwise healthy individuals (e.g. thrush), associated with high morbidity rates (5). In specific clinical settings where patients are subjected to immunosuppressive treatments, complex medical procedures or, as recently seen in cases of underlying infection (e.g., COVID-19 (6)), these superficial infections may lead to life-threatening systemic candidiasis through the translocation of the fungus to the bloodstream. These candidaemias are the fourth most common nosocomial bloodstream infections and have associated mortality rates of 45-75% (2).

Epithelial cells are the first barrier encountered by microbes in mucosal tissues and are in continuous contact with the millions of microbes that constitute the host microbiota. Therefore, epithelial discrimination between commensal and pathogenic microbes in collaboration with resident immune cells in mucosal tissues of healthy individuals is essential for maintaining homeostasis and proper immune responses to clear pathogens. In the oral cavity, interactions between epithelial cells and *C. albicans* are greatly influenced by morphological switching of the fungus, with the yeast form being associated with commensal colonisation and hyphae with invasion and pathogenesis (7). Activation of epithelial cells by *C. albicans* yeasts promotes the induction of phosphoinositol-3-kinase/protein kinase B (PI3K/Akt) and nuclear factor kappa-light-chain-enhancer of activated B cells (NF-κB) signalling pathways, and a weaker, transient activation of the mitogen-activated protein kinase (MAPK) signalling pathways; p38, c-Jun N-terminal kinases (JNK), and extracellular signal-regulated kinases (ERK1/2). In contrast, hyphal growth results in the strong, sustained activation of all three MAPK signalling pathways in addition to the continued activation of NF-κB and PI3K/Akt. This leads to the recruitment of the AP-1 transcription factor component c-Fos, and expression of a plethora of cytokines, chemokines and antimicrobial peptides (8, 9). This hypha-driven signalling circuit is triggered by secretion of the toxic peptide candidalysin from growing hyphae (10, 11), detected through a molecular circuit dependent on EGFR (12, 13). This cytolysin is the key virulence factor in inducing cell damage and immune activation in epithelial cells during *C. albicans* infections (14).

Even though our understanding of the molecular interplay between *C. albicans* and epithelial cells has greatly improved in the last decade, many aspects of the early events occurring during these interactions are as-yet unknown. In particular, the transcriptional regulation machinery activated in both invasion and commensalism remain to be fully uncovered. Here, we identify early growth response protein 1 (EGR1) as an important transcription factor expressed in human oral epithelial cells (OECs) in response to *C. albicans*. Importantly, we show that OECs increase EGR1 expression in response to *C. albicans* in a manner independent of fungal morphology, viability or candidalysin production. Therefore, this transcription factor is not only involved in the response to filamentous growth, but also mediates epithelial cell activation by yeasts. We show that EGR1 expression induction by *C. albicans* occurs through Raf, ERK1/2 and NF-κB, but not via PI3K/mTOR, and that this transcription factor potentiates the immune response of OECs to the fungal presence by regulating pro-inflammatory cytokine release.

## RESULTS

### Fungal sensing by oral epithelial cells activates the expression of EGR1

Previous work from our group identified activation of NF-κB and AP-1 family transcription factors in OECs in response to infection by *C. albicans* (8, 15, 16). In addition, analysis of the OEC transcriptome identified several key transcription factors that were upregulated in *C. albicans*-infected reconstituted oral epithelium (ROE) at 6 h and 24 h post-infection (15). Among the most significantly upregulated genes were two members of the early growth response (EGR) family; EGR1 and EGR3 (Table 1). To validate the results of this microarray analysis, expression of these two genes was determined over the first 4 h of infection in TR146 OEC monolayers using RT-qPCR (Figure 1A). *EGR1* expression significantly increased 18.5-fold (*P* < 0.01) at 2 h, peaking at 20.1-fold (*P* < 0.01) by 4 h post-infection, whilst *EGR3* gene expression increased 15.8-fold (*P* < 0.05) at 2 h, increasing further to 32.5-fold (*P* < 0.0001) by 4 h post-infection. However, only EGR1 showed significant increases in protein levels in infected cells (Figure 1B) with significantly elevated protein levels at 2 h (2.2-fold; *P* < 0.01) and 4 h (2.3-fold; *P* < 0.01). Even though the gene expression of *EGR3* was greater than *EGR1* in TR146 monolayers, there was no corresponding increase in EGR3 protein production at the time points observed.

**Table 1.** EGR family members are expressed following *Candida albicans* infection of reconstituted oral epithelium. Microarray expression data of EGR family members whose expression was significantly different after *C. albicans* infection in reconstituted human oral epithelium (ROE).

| Gene symbol | 6h post-infection |  | 24h post-infection |  |
| --- | --- | --- | --- | --- |
|  | Fold change<br>(*Candida* vs. PBS) | *P*-value<br>(*Candida* vs. PBS) | Fold change<br>(*Candida* vs. PBS) | *P*-value<br>(*Candida* vs. PBS) |
| EGR1 | 4.0 | 4.61 | 5.3 | 3.38 |
| EGR3 | 4.3 | 7.03 | 32.4 | 2.75 |

**Figure 1.**
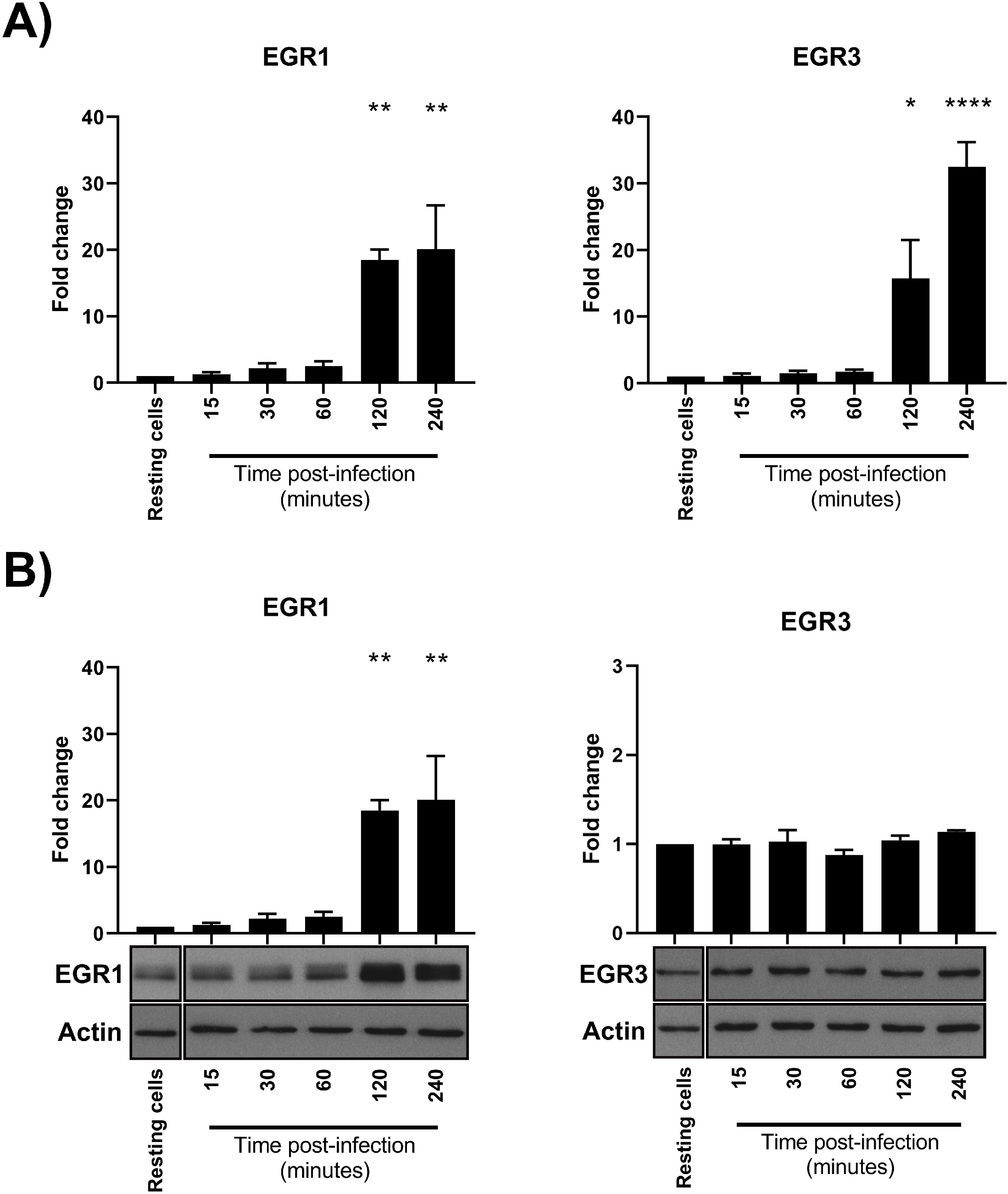
EGR1 is expressed following *Candida albicans* infection of oral epithelial cells. EGR family members whose expression was significantly different after *C. albicans* infection of reconstituted human oral epithelium (ROE) via microarray analysis were studied in TR146 cell monolayers by qRT-PCR (A) Western blot (B). EGR1 overexpression was observed at both RNA and protein levels, whilst only RNA upregulation was detected for EGR3. All data is shown as mean fold change ± S.E.M. Significance calculated relative to resting cells, \**P*<0.05, \*\**P*<0.01.

### Induction of EGR1 is independent of *C. albicans* major virulence factors

A key feature of *C. albicans* pathogenesis and hallmark of the switch from commensal to invasive growth is the formation of filamentous hyphae (17). To determine whether hyphal growth was required to induce *EGR1* gene expression, TR146 monolayers were infected with the yeast-locked *efg1*Δ/Δ/*cph1*Δ/Δ null mutant or wild-type BWP17+cIP30 *C. albicans* strains. Infection with both strains resulted in a significant increase in expression of *EGR1* as early as 15 min post-infection (Figure 2A). However, this increased *EGR1* expression was significantly greater at 2 and 4 h post-infection with BWP17+cIP30. Crucially, the protein levels of EGR1 produced during infection with both strains followed the same profile and were significantly greater than in resting cells at 2 h post—infection, and not significantly different between strains (Figure 2B). This suggests that the production of EGR1 in the early stages of infection is not dependent on fungal morphology.

**Figure 2.**
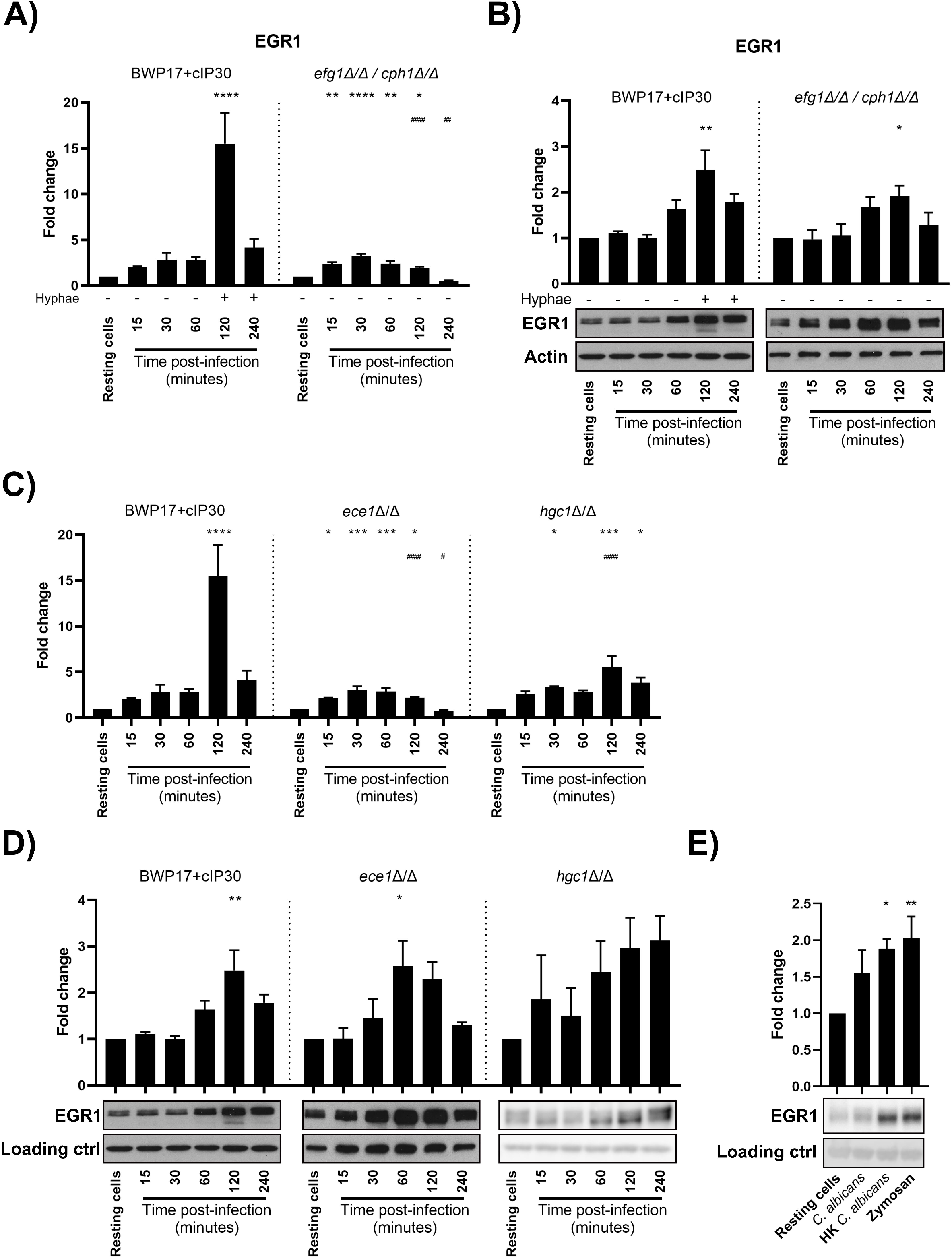
Production of EGR1 is independent of major fungal virulence factors. EGR1 RNA (A) and protein (B) levels in TR146 epithelial cell monolayers following infection with a parental strain of *C. albicans* BWP17+cIP30 or the yeast-locked mutant *efg1*Δ/Δ/*cph1*Δ/Δ. EGR1 RNA (C) and protein (D) levels in TR146 cells post-infection with *C. albicans* BWP17+cIP30, a Candidalysin-deficient *ece1*Δ/Δ strain, or an *hgc1*Δ/Δ mutant, which is unable to sustain hyphal growth. (E) Western blot analysis of EGR1 in TR146 cells treated with live wild-type *C. albicans*, heat-killed yeast, or fungal cell wall components (zymosan, 50 μg/mL; chitin, 2 μg/mL; *N*-mannan, 50 μg/mL; *O*-mannan, 25 μg/mL) for 2 h. All data is shown as mean fold change ± S.E.M. Significance calculated using one-way ANOVA: \**P*<0.05, \*\**P*<0.01, \*\*\**P*<0.001, \*\*\*\**P*<0.0001, relative to vehicle control; #*P*<0.05, ##*P*<0.01, ####*P*<0.0001, relative to parental strain.

Although hyphal growth is required to cause damage to epithelial cells, it is the peptide toxin candidalysin that causes the cellular damage, and not the hyphae (10). Candidalysin is encoded by the *ECE1* gene, which is highly expressed in hyphae. In common with other microbial pore forming toxins, this toxin can induce epithelial intracellular signalling and gene transcription at sub-lytic concentrations (14). The contribution of candidalysin to the production of EGR1 was investigated using an *ECE1* gene deficient strain (*ece1*Δ/Δ) and an *hgc1*Δ/Δ mutant, which expresses *ECE1* at wild-type levels (10, 18), but cannot sustain hyphal growth. Although there was reduced expression of *EGR1* in response to *ece1*Δ/Δ and a *hgc1*Δ/Δ infected cells relative to the wild-type infection, there was still a significant increase in *EGR1* expression relative to resting cells (Figure 2C). Additionally, despite a slight reduction in gene expression levels, EGR1 production appears to be independent of candidalysin as there was no significant difference in protein levels following infection by *ece1*Δ/Δ or either of the toxin-producing strains (Figure 2D). The presence or absence of hyphae also had no significant effect on EGR1 protein levels.

As EGR1 is produced by *C. albicans* infected cells in a manner independent of hyphal growth and candidalysin production, we next determined whether fungal viability was required. OECs were infected with live *C. albicans* or treated with heat-killed (HK) *C. albicans* yeast or cell wall components for 2 h. HK *C. albicans* elicited production of EGR1 to a similar degree as live infection, as did empty fungal cell walls; the Dectin-1 agonist zymosan (Figure 2E).

### The MAPK ERK1/2 and EGFR is required for EGR1 production in response to live *C. albicans* and fungal components

As *C. albicans* can colonise the oral cavity as a commensal or a pathogen, it is critical for the host to discriminate between each state appropriately. Previously, we identified that NF-κB and PI3K/mTOR signalling were both involved in epithelial cell recognition of yeast morphotypes and low fungal burdens. However, a second, strong MAPK response was required to discriminate commensal yeast from the invasive hyphae (8, 15). To determine which of these pathways were involved in eliciting the production of EGR1, MAPKs, PI3K/mTOR and NF-κB activation were inhibited using small chemical inhibitors, and EGR1 protein levels quantified 2 h post-infection (Figure 3A). EGR1 levels were significantly diminished following inhibition of NF-κB (*P* < 0.01) and the MAPK kinase kinase (MAP3K), Raf1 (*P* < 0.001), with EGR1 levels being reduced to less than that of resting cells. Interestingly, of the MAPKs, only ERK1/2 inhibition significantly reduced EGR1 production (*P* < 0.001) with inhibition of JNK or p38 having no significant effect. To test whether the production of EGR1 was dependent on ERK1/2 in response to other stimuli, OECs were treated with HK *C. albicans* or zymosan for 2 h following ERK1/2 inhibition (Figure 3B). Western blot analysis revealed that when ERK1/2 phosphorylation and signalling was supressed, EGR1 production was abolished in response to all the stimuli tested.

**Figure 3.**
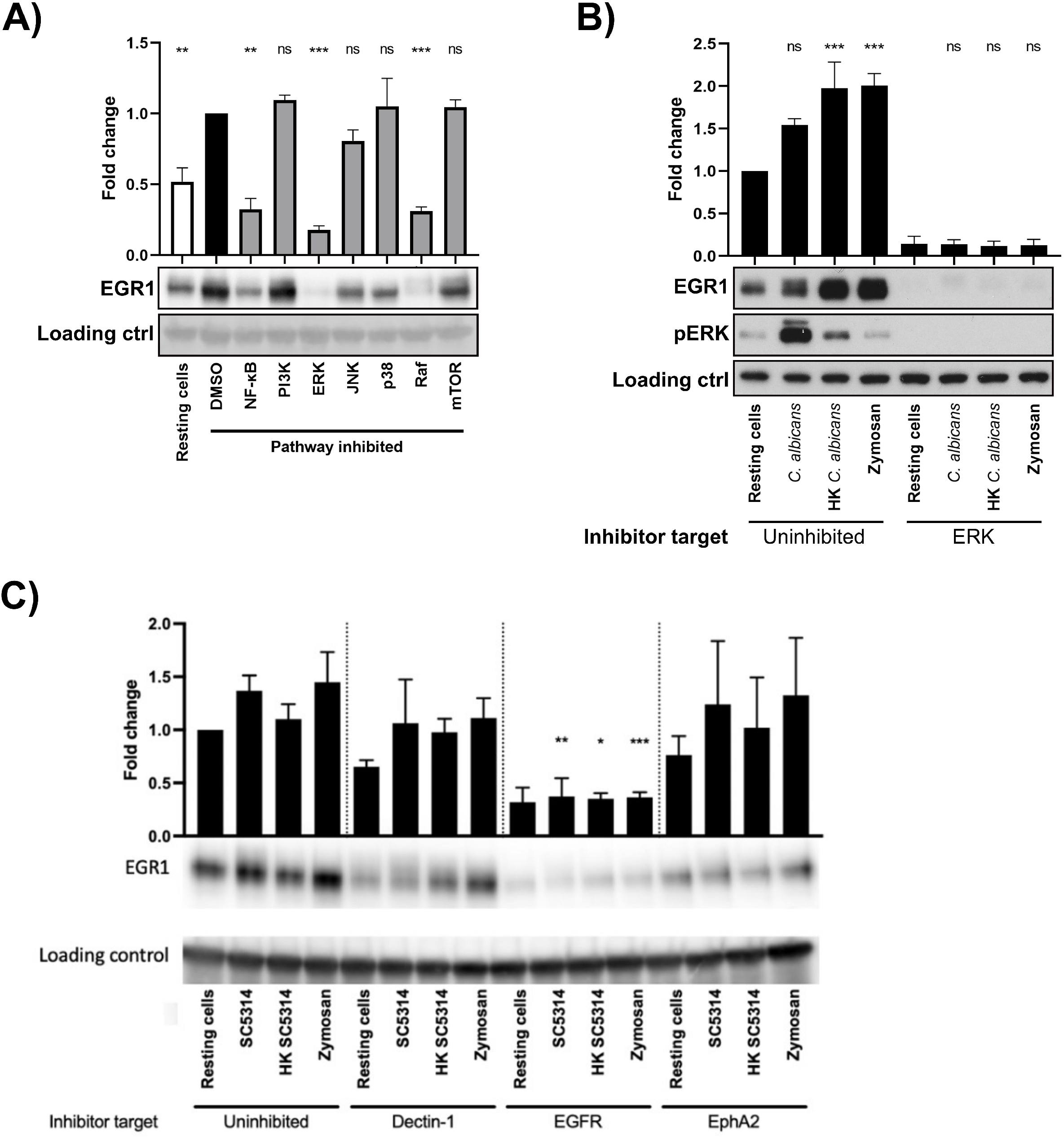
NF-κB, ERK1/2, Raf, and EGFR signalling control EGR1 increase upon fungal infection. **(A)** Western blot analysis of EGR1 in TR146 epithelial cell monolayers. Cells were treated with inhibitors targeting components of signalling pathways, or a vehicle control of DMSO, prior to infection with *C. albicans*. Significance calculated relative to vehicle control (DMSO). **(B)** EGR1 protein levels in cells stimulated with live or heat-killed (HK) *C. albicans* or zymosan for 2 h after treatment with an ERK1/2 inhibitor or a vehicle control of DMSO. **(C)** Dectin-1, EGFR, and EphA2 were inhibited with laminarin, gefitinib, and dasatinib, respectively, and then cells were stimulated with live or heat-killed (HK) *C. albicans* or zymosan (50 μg/mL) for 2 h. Western blotting was then used to measure EGR1 expression. Significance calculated relative to resting cell controls. All data is shown as mean fold change ± S.E.M. Significance calculated using one-way ANOVA: \**P*<0.05, \*\**P*<0.01, \*\*\**P*<0.001.

The recognition of microbe-associated molecular patterns is usually mediated by pattern recognition receptors (PRRs). Several PRRs are involved in the recognition of *C. albicans* by OECs; the Ephrin A2 receptor (EphA2) and Dectin-1 sense β-glucans on the fungal cell surface (19). Fungal invasins bind E-cadherin to induce endocytosis (20), which is enhanced by HER2 and EGFR (21). EGFR is also indirectly activated by candidalysin (12), leading to activation of MAPK signalling and expression of cytokines (13). Given this, the role of Dectin-1, EGFR and EphA2 in the induction of EGR1 expression were investigated using inhibitors (Figure 3C). Following inhibition of EGFR, EGR1 levels were significantly deceased in response to infection with *C. albicans* (*P* < 0.01), treatment with HK yeast (*P* < 0.05) and zymosan (*P* < 0.001) after 2 h. Although there was not a significant decrease, EGR1 levels appeared to be slightly diminished when EphA2 was inhibited.

### EGR1 regulates expression of cytokines following recognition of *C. albicans*

The responses of epithelial cells are critical for generating and maintaining immune tolerance, as well as alerting the immune system and promoting an inflammatory response to pathogens. To investigate whether EGR1 plays a protective role in host responses to *C. albicans*, the impact of knock down of *EGR1* mRNA on cellular damage and cytokine production was determined using siRNAs. OECs were transfected with siRNAs (either a single target, or a pool of siRNAs) to knockdown EGR1 expression. Both single and pooled siRNAs were able to abolish EGR1 production in response to *C. albicans* infection (Figure 4A). Infection of OECs treated with EGR1 siRNA for 24 h had no impact on fungal-induced OEC cellular damage as measured by LDH release (Figure 4B). There was, however, a significant decrease in the secretion levels of GM-CSF, IL-1α and IL-1β. Interestingly, the levels of IL-6 and IL-8 were increased following knock-down of EGR1 (Figure 4C).

**Figure 4.**
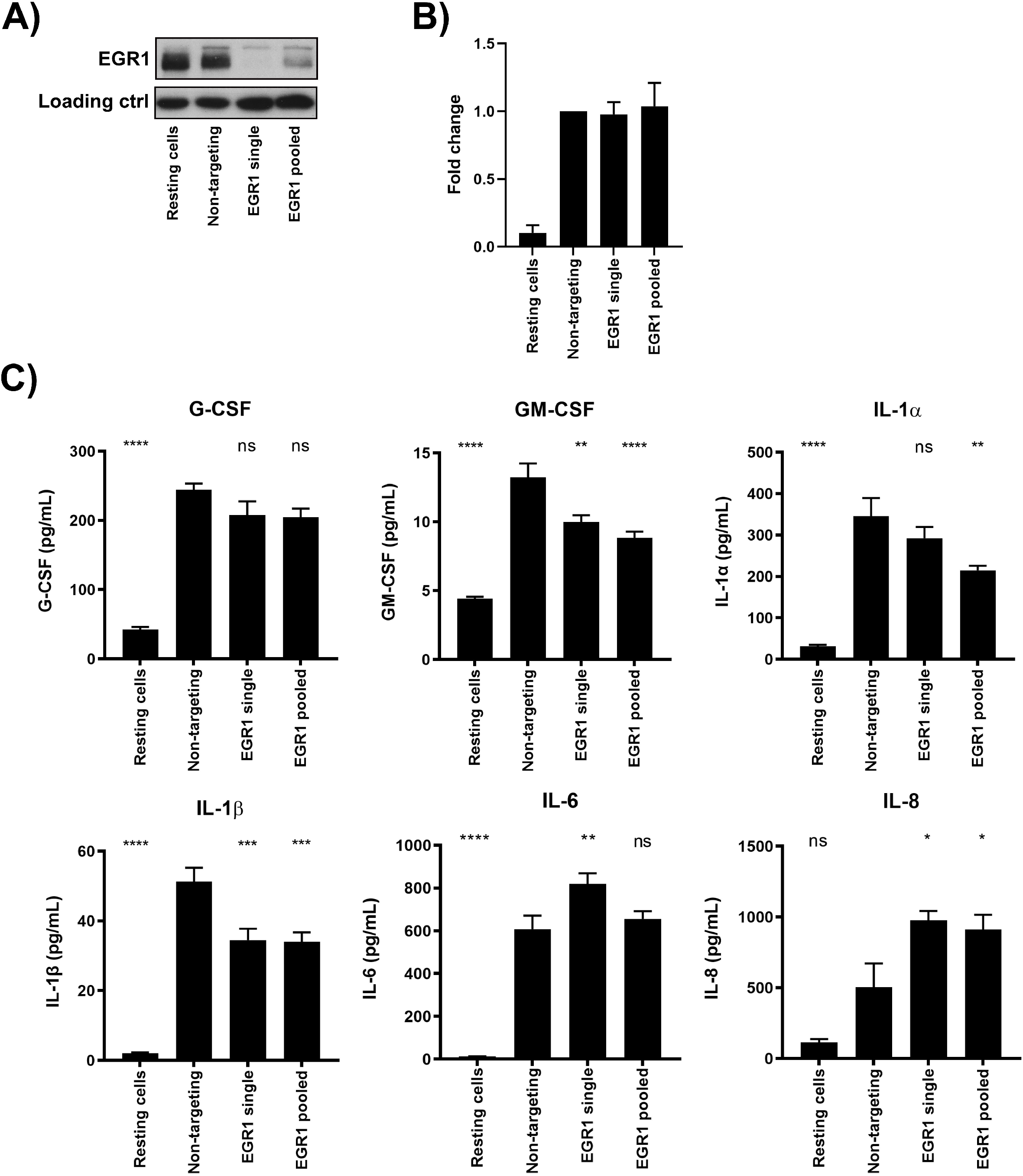
EGR1 regulates the production of cytokines in oral epithelial cells. TR146 cells were treated with a single siRNA or a pool of siRNAs targeting EGR1 and then infected with *C. albicans*. **(A)** Western blot analysis to confirm knockdown of EGR1 in OECs. **(B)** LDH quantification and **(C)** Luminex analysis of culture supernatant at 24 h post-infection. All data is shown as mean fold change ± S.E.M. Significance relative to non-targeting siRNA control calculated using one-way ANOVA: \**P*<0.05, \*\**P*<0.01, \*\*\**P*<0.001.

## DISCUSSION

Epithelial cells are one of the first host cells to encounter microbes, including potential pathogens, and orchestrate the immune responses that maintain homeostasis. As *C. albicans* colonises mucosal surfaces as both a commensal and pathogen, it is crucial for the immune system to be able to identify and respond to switches between these states. A biphasic MAPK response allows epithelial cells to discriminate between the commensal and pathogen states and involves an initial morphology-independent response, followed by a second hyphal-specific activation dependent on fungal burden that alerts the host to infection (8, 11). Utilising previously published microarray data (15), we sought to understand the host responses to early infection by this opportunistic pathogen. Here we identify EGR1 induction as a morphology-, viability- and virulence-independent response that alerts the host to the presence of *C. albicans*, without inducing a potent proinflammatory response. The data confirms that epithelial cells recognise and specifically respond to both colonising and invasive fungal states.

EGR1 is a zinc-finger transcription factor expressed by different cell types and induced in response to various stimuli, including physical, chemical and biological factors (22–25). However, little is known about its relevance in epithelial responses to microbes, with only a few studies analysing EGR1 induction upon infection with bacteria (24, 26) or the fungal pathogen *Aspergillus fumigatus* (27). The degree of EGR1 expression in response to bacterial species is dependent on host cell type and bacterial viability, as dead bacteria had a diminished capacity or were incapable of eliciting EGR1 expression (24). Our data shows that EGR1 is expressed by epithelial cells within two hours of exposure to *C. albicans* to both yeast-locked and hyphal-competent strains. Furthermore, EGR1 is produced in response to heat-killed yeast and the cell wall component zymosan, indicating that the EGR1 response is independent of morphology and viability. These data indicate that EGR1 induction is a feature of epithelial recognition of *Candida* presence and is not involved in the discrimination between commensal and pathogenic states of *C. albicans*.

In the early stages of infection with *C. albicans*, the morphology-independent activation of MAPKs coincides with NF-κB and PI3K activation. We demonstrate that ERK1/2, NF-κB and the MAP3K Raf1 were indispensable for EGR1 induction during infection. In the above-mentioned studies, *C. albicans* infection leads to the expression of EGR1 via the MAPKs ERK1/2, p38 or JNK, or a combination of all three. The upstream signalling events that lead to activation of these factors is unclear, so the finding that Raf1 plays a role in EGR1 induction, in addition to ERK1/2 signalling, is a notable finding. Our data demonstrate a Raf1-MEKK1/2-ERK1/2-EGR1 cascade that leads to activation of cytokine production. Although the three MAPK signalling pathways are essential in host epithelial cell responses to *C. albicans*, the microbial triggers have not been well described. In the case of *C. albicans*, ERK1/2 mediates the production of EGR1 in response to live and HK yeast, as well as zymosan, suggesting that the signalling cascades triggered converge on ERK1/2. Furthermore, the key role we identified for NF-κB signalling in inducing EGR1 production shows that EGR1 is regulated by general fungal recognition (8, 15), rather than invasive infection (8). The data indicates that this initial recognition interaction between epithelial cells and *C. albicans* may result in a “priming” event that potentiates future production of cytokines and damage responses resulting from hyphal invasion, given that inhibition of cytokine production is only partial. Thus, initial recognition of the yeast form of the fungus might lead to the OECs being primed to respond more strongly to subsequent hyphal growth in a process akin to innate immune training.

Induction of innate immune responses to fungal cells is triggered by interaction of specific receptors via several fungal-specific microbial associated molecular patterns. Key in these interactions are those between the fungal cell wall component, β-glucan and the C-type lectin receptor, Dectin-1 (28). Although zymosan is a ligand of Dectin-1, the inhibition of Dectin-1 did not significantly alter the levels of EGR1 during treatment with zymosan, or during infection with *C. albicans*. This is not unexpected, as we have previously shown that *C. albicans* infection of OECs does not lead to the phosphorylation of Syk (8), a key component of the canonical Dectin-1 signalling pathway. In contrast to immune cells, we have previously demonstrated that the fungal peptide toxin candidalysin is critical for *C. albicans* virulence at mucosal surfaces, as well as recognition of invasive hyphae (10). This cytolysin is produced during hyphal growth and activates MAPK signalling indirectly via the EGFR receptor, leading to cytokine release (10, 12). The signalling cascade activated by EGFR is dependent on the adapters Gab1, Grb2 and Shp2 (13), and different cytokines are produced depending on the MAPK activated (29). Utilising an *ECE1* gene deficient strain of *C. albicans*, we determined that the induction of EGR1 was not dependent on candidalysin. However, EGFR inhibition significantly reduced EGR1 production in response to wild-type *C. albicans* infection. Lying upstream of ERK1/2, Raf1 is phosphorylated by Ras-GTPase, which in turn can be activated by the EGFR receptor via the adapters Grb2 and Son of Sevenless (SOS) (30, 31). In addition to Dectin-1, fungal β-glucan can also be detected via EphA2 (19). Notably, EGFR can form complexes with EphA2 (32) or HER2 (21) to endocytose fungal hyphae via the fungal adhesin/invasin Als3. As such, EphA2 may also be required for the induction of EGR1 in OECs.

Although EGFR activation leads to cytokine production following activation by candidalysin, the cytokine release induced by the biphasic MAPK response appears to be driven by the hyphal-specific secondary phase, rather than the initial morphology-independent activation. Although the depletion of EGR1 led to a significant decrease in the levels of GM-CSF, IL-1α and IL-1β, there was no change in the production of G-CSF and an increase in IL-6 and IL-8. Further, knockdown of EGR1 did not alter LDH levels in culture supernatant, which is an indicator of cell damage. This indicates that although EGR1 can induce the expression of cytokines, it does not play a protective role against the damage caused during *C. albicans* infection.

In summary, our findings demonstrate that EGR1 induction plays a role in the control and regulation of early responses to fungal contact in the oral epithelium via the activation of EGFR-MAPK signalling, independently of morphology and candidalysin, two major virulence factors of *C. albicans*. As such, EGR1 likely plays a role in host homeostasis by driving a priming response to better respond to a possible invasive infection, rather than inciting a proinflammatory response.

## MATERIALS AND METHODS

### Cell Lines, Reagents, and *Candida* Strains

Experiments were carried out using the TR146 buccal epithelial carcinoma cell line. Monolayer cultures were grown in DMEM/F-12 with 15% FBS, and experiments carried out in serum-free media.

Inhibitors of ERK1/2 (PD184352; 2μM) and Dectin-1 (laminarin, 1mg/mL) were purchased from Sigma. Inhibitors of JNK (SP600125; 10 μM), p38 (SB203580; 10 μM), mTOR (Ku-63794; 10 μM), NF-κB (BAY11-7082; 2 μM) and RAF1 (RAF1 Kinase inhibitor I; 10 μM) were from Calbiochem. The inhibitor of EGFR (gefitinib, 1 μM) was from SelleckChem and JNK (SP600125, 10 μM) was from Cell Guidance Systems. Primary antibodies to EGR1, EGR3, phospho-ERK1/2, GAPDH and actin, and an inhibitor of EphA2 (dasatinib, 100 nM) were from Cell Signaling Technologies. Secondary HRP-conjugated secondary antibodies used were goat anti-rabbit (Jackson ImmunoResearch) and goat anti-mouse (Merck).

Fungal strains used in this study were *C. albicans* wild-type SC5314 (33), *efg1*Δ/Δ/*cph1*Δ/Δ (34), BWP17+CIp30 (35), *hgc1*Δ/Δ (36), *hgc1*Δ/Δ+*HGC1* (36), *ece1*Δ/Δ (10), *ece1*Δ/Δ+*ECE1* (10). Heat-killed SC5314 was prepared from overnight cultures via incubation at 57°C for 1 h. Cell monolayers were infected with live or heat-killed fungus at a multiplicity of infection (MOI) of 10. Cell wall components purified from *C. albicans* cell walls; zymosan (50 μg/mL), chitin (2 μg/mL), *N*-mannan (50 μg/mL) and *O*-mannan (25 μg/mL), were kind gifts from Neil Gow.

### RNA Extraction and Analysis

RNA was isolated from oral epithelial cells using the Nucleospin II kit (Macherey-Nagel) and treated with Turbo DNase (Invitrogen) to remove genomic DNA contamination. cDNA was generated using SuperScript IV Reverse Transcriptase (Invitrogen) and real-time PCR was performed using Jumpstart SYBR green mastermix (Sigma-Aldrich) on a Rotorgene 6000 (Corbett Research). EGR1 (forward 5’-GCCTGCGACATCTGTGGAA-3’; reverse 5’-GCCGCAAGTGGATCTTGGTA-3’), EGR3 (forward 5’- CCATGATTCCTGACTACAACCTC-3’; reverse 5’-GTCTTTGAATGCCTTGATGGTCT-3’), and YWHAZ (forward 5’-ACTTTTGGTACATTGTGGCTTCAA-3’; reverse 5’-CCGCCAGGACAAACCAGTAT-3’) primers were taken from Primerbank (http://pga.mgh.harvard.edu/primerbank/) (37).

Transcriptional data from three-dimensional models of reconstituted oral epithelium (ROE) previously published by our group (15) was used to analyse the impact of *C. albicans* infection on the gene expression of EGR gene family. Data was statistically analysed using Partek Genomics Suite (version 6.4) and significance, relative to sham-infected controls, calculated using one-way ANOVA with false discovery rate (FDR) correction.

### Western Blotting

Epithelial cells were lysed using a modified RIPA lysis buffer containing protease and phosphatase inhibitors (Sigma-Aldrich). Lysates were incubated on ice for 30 min and centrifuged for 10 min at 4°C. Supernatants were assayed for total protein using the BCA protein quantitation kit (Thermo Fisher Scientific). 20 μg of protein was separated on 12% ProtoGel Quick-Cast Gels (National Diagnostics) before transfer to nitrocellulose membranes (BioRad). After probing with primary and secondary antibodies, membranes were developed using Immobilon chemiluminescent substrate (Millipore) and exposed to ECL film (GE Healthcare) or luminescence captured with an ImageQuant 800 (Amersham). Densitometry analysis of Western blots was performed using Fiji (Version 2.3.0) (38).

### RNA Interference

Knockdown of gene expression was performed using previously reported siRNAs for EGR1 (39), or commercially available siRNA pools for EGR1 and EphA2 (Santa-Cruz Biotech). As a control, a non-silencing siRNA duplex with no homology to known human genes was used (Santa-Cruz Biotech). Transfection of siRNA was performed overnight using 6 nM for each siRNA with HiPerFect reagent (QIAGEN) according to the manufacturer’s reverse transfection protocol. Protein analysis demonstrated that transcript levels for targeted genes were reduced and maintained this level for at least a further 24 h.

### Analysis of cell damage and cytokine Determination

Lactate dehydrogenase (LDH) in culture supernatant was quantified using the Cytotox-96 non-radioactive cytotoxicity assay (Promega). Cytokine levels in cell culture supernatants was determined using the Magnetic Luminex Performance Assay (Human Base Kit A; R&D Systems, coupled with the Bio-Plex 200 (Bio-Rad). The trimmed median value was used to derive the standard curve and calculate sample concentrations.

### Statistics

Statistical analyses of data (except microarray data) were performed using GraphPad Prism 7 (version 7.0b; GraphPad Software Inc.) and significance calculated with one-way ANOVA; *p<0.05, **p<0.01 ***p<0.001, ****p<0.0001.

## Acknowledgements

DLM was supported by the Biotechnology Biological Sciences Research Counci (BBSRC) (BB/S016899/1, BB/M009513/1). JRN was supported by grants from the Wellcome Trust (214229_Z_18_Z), and the National Institute for Health and Care Research (NIHR) Biomedical Research Centre based at Guy’s and St Thomas’ NHS Foundation Trust and King’s College London and/or the NIHR Clinical Research Facility. DLM and JRN were supported by the National Institutes of Health (DE022550).

